# Sustained cross-species transmission of gammacoronavirus in wild birds reveled by viral characterization in China

**DOI:** 10.1101/2025.04.17.648926

**Authors:** Dan-Shu Wang, Ying-Ying Zou, Qian Liu, Hao Huang, Pei-Yu Han, Jun-Ying Zhao, Li-Dong Zong, Ye Qiu, Yun-Zhi Zhang, Xing-Yi Ge

**Author notes:** Correspondence (Y.-Z.Z.); (X.- Y.G.).

## Abstract

Gammacoronavirus (γ-CoV) primarily infects poultry, wild birds, and marine mammals. The widespread distribution and circulation of γ-CoV in the ecological environment may lead to sustained transmission and economic loss. To better understand the diversity of γ-CoV in wild birds, we collect 482 wild-bird fecal samples from Yunnan, encompassing fourteen bird species. We detected twelve γ-CoV positive samples in five bird species, with the characterization of five complete genomes - HNU5-1, HNU5-2, HNU5-3, HNU6-1, and HNU6-2-indicating that these genomes represent two viral species. The HNU5 strains were derived from Black-headed gull (*Chroicocephalus ridibundus*), while the HNU6 strains were came from Mallard (*Anas platyrhynchos*), and both of those were recombinant. The HNU5 strain exhibited the highest sequence identity (95.45%) with a γ-CoV strain isolated from *Numenius phaeopus* (GenBank accession: PP845452). Similarly, the HNU6 strain showed 95.18% nucleotide identity with a γ-CoV strain (GenBank accession: PP845437) derived from *Anas platyrhynchos*. Taxonomic analysis confirmed that HNU6s belong to the *Gammacoronavirus anatis* species, while HNU5s attributed to a new species. Cross-species analysis revealed active host-switching events among γ-CoVs, indicating potential transmission of γ-CoVs from marine mammals to wild bird, from wild bird to poultry, and inter-wild bird and inter-poultry transmission. In summary, we report five new γ-CoV strains in wild birds and outline the cross-species transmission of γ-CoVs. Our findings link γ-CoV hosts across different natural environments and provide new insights for exploring γ-CoVs.

## 1 Introduction

Coronaviruses (CoVs), which belong to the *Nidovirales* order, *Coronaviridae* family, and *Orthocoronavirinae* subfamily, are large, enveloped, single-stranded, positive-sense RNA viruses [1, 2]. The International Committee on Taxonomy of Viruses (ICTV) systematically divides CoVs into four genera: *Alphacoronavirus* (α-CoV), *Betacoronavirus* (β-CoV), *Deltacoronavirus* (δ-CoV), and *Gammacoronavirus* (γ-CoV). The γ-CoV includes three subgenera: *Brangacovirus*, *Cegacovirus* and *Igacovirus*. There are five verified species within the γ-CoV genus: (i) *Gammacoronavirus galli*, which includes several genotypes of infectious bronchitis virus (IBV), (ii) *Gammacoronavirus pulli*, comprising IBV and Turkey CoVs, (iii) *Gammacoronavirus anatis* (DuCoV_2714), which includes duck CoVs, (iv) *Gammacoronavirus brantae* (BcanCoV_CB17), and (v) *Gammacoronavirus delphinapteri*, which includes beluga whale coronavirus (BWCoV) and bottlenose dolphin coronavirus HKU22 (BdCoV_HKU22).

CoVs infect mammals and birds, with members of α-CoVs and β-CoVs only infect mammals, such as bats, mice, and humans [3]. Traditionally, δ-CoVs and γ-CoVs predominantly infect birds, indicating that birds are the natural reservoirs for these CoVs. Since the first avian coronavirus, infectious bronchitis virus (IBV), was described in 1936, δ-CoVs and γ-CoVs have been successively detected in other wild bird species [4–7]. To date, wild birds across fifteen avian orders and thirty families, comprising 108 species, have been found to carry δ-CoVs and γ-CoVs [8]. The Anseriformes and Charadriiformes orders are the primary wild bird orders that harbor CoVs, with γ-CoVs being the most commonly found type, followed by δ-CoVs [9].

In addition to wild birds, δ-CoVs and γ-CoVs can also infect mammals. The Porcine deltacoronavirus (PDCoV) of δ-CoV were detected in domestic pigs in China the Unites states, Canada, and may other countries [10, 11] , causing severe diarrhea and vomiting[4]. Additionally, a novel γ-CoV was identified in 2008 from a liver sample of a beluga whale in the USA, making the first discovery of γ-CoV in a mammal[12]. Recently, γ-CoVs have also been detected in fecal samples from bottlenose dolphins [13]. Although γ-CoVs are typically found in captive cetaceans, a strain was detected in a wild striped dolphin in 2024 [14], indicating γ-CoV may be widely prevalent in marine wildlife cetaceans.

At present, there are more than 9,000 known bird species distributed across various geographic regions. Based on their habitat, birds can be categorized into two groups: poultry and wild birds. Poultry, as an important economic livestock, is widely distributed around the globe, while wild birds possess not only ornamental value but also ecological significance. Wild birds exhibit unique wintering behaviors. Intensive flight activities can lead to immunosuppression, making wild-birds more susceptible to pathogens [15]. Furthermore, pathogens carried by wild birds may adapt and jump to some mammalian species, including humans, due to long-distance migration [16]. It is reported that 1.4% of viruses transmitted from nonhumans to humans are found in Anseriformes [17]. A notable case of wild birds transmitting a virus to humans is the H5N1 influenza virus [18]. As the natural reservoir of γ-CoV, birds can also can transmit γ-CoV though long-distance flying. The highly variability and mutation rate of CoVs enable them to conduct host-switching [19, 20]. IBV-like strains have been detected in wild birds in both Brazil and Australia [21, 22], while fecal samples of domestic ducks and geese have shown the presence of H120-like IBV [23], indicating that γ-CoV can transmit from poultry to wild birds and vice versa.

To mitigate the risk of avian-to-human transmission, it is crucial to monitor the prevalence of viruses in birds. Yunnan province, located in the southwestern part of China, serves as a natural ecological cradle that attracts a large number of wild birds for breeding. Statistically, Yunnan is home to over 1,000 bird species, accounting for about 9% of the world’s total. Previously, we detected a series of new δ-CoVs from wild birds in Yunnan indicating the widespread distribution of CoVs in the region [7]. The spread of γ-CoVs may impact the health of bird populations and ecological balance, making it vital to conduct ongoing monitoring and research on γ-CoVs.

This study aims to detect and characterize γ-CoVs in wild birds and explore their genetic diversity and evolution, and the co-evolution between γ-CoVs and their hosts, thereby revealing host-switching events of γ-CoVs among hosts in different ecological environment.

## 2 MATERIALS AND ETHODS

### 2.1 Sample collection and RNA extraction

Fecal samples were collected in Kunming and Dali, Yunnan, China in Dec 2021 and Mar 2022, respectively. Samples were put into Viral Transport Media (VTM), and transported to the lab with the dry ice, stored in the environment of -80℃. Then, virus RNA of the fecal samples was extracted with the TIANamp Virus DNA/RNA Kit (DP315, TIANGEN), following the manual. RNA was eluted into 60μl RNase-free water and stored in the environment of -80℃ waiting for detection.

### 2.2 CoV detection and DNA sequencing

To screening γ-CoV, pairs of specific primers were synthesized which aims to the RNA-dependent RNA polymerase (*RdRp*) gene (Table S1) of γ-CoV [24]. The target fragment size of the RT-PCR of γ-CoV is 440bp. PCR amplification was performed using the PrimeScript™ One Step RT-PCR Kit Ver.2 (Takara). The PCR mixture (20 μL) contained 2 μL of extracted RNA, 12.5 μL 2 × 1 Step Buffer and 1 μL PrimeScript 1 Step Enzyme Mix. The mixtures were denaturation in 50℃ for 30 min firstly, and amplified by 40 cycles of 94℃ for 2 min, 94℃ for 30 s, 55-65℃ for 30 s, and 72℃ for 1 min/kb and a final extension at 72℃ for 10 min. To confirm the bird species, the 12S rRNA gene and 16S rRNA gene (Table S1) were amplified and sequenced [25].

All products that glowed at the corresponding position were gel-purified and sequenced. Then, the PCR product sequences were compared with gene sequences in GenBank.

### 2.3 Complete genome sequencing

Five γ-CoVs complete genomes were amplified and sequenced using the PrimeScript™ One Step RT-PCR Kit Ver.2 (Takara). The first-round fragment was amplified using degenerate primers based on multiple alignments of complete CoV genomes from GenBank. Additional primers were derived from the results of the first and subsequent rounds of sequencing results. The 5′ and 3′ ends sequences were obtained by 5′ and 3′ RACE (Roche). Sequences were assembled to obtain the full-length genome sequences.

### 2.4 Genome analysis

The annotation of genome was integrated by using ORFfinder and Corsid [26], which the former provides the putative open reading frames (ORFs) and the latter offer the predicted transcription regulatory sequences (TRS). The annotated gene was compared to other CoVs using BLAST to ensure the ORFs is named correctly. The nucleotide sequence identity was analyzed by BioAider v1.527 [27]. Sequences alignment was executed by using MAFFT v7.149 program in BioAider v1.527 [27, 28]. A comprehensive search of the GenBank database using the taxonomic identifier “taxid 694013” retrieved 19,085 γ-CoV sequences. These sequences were utilized as query sequences against a reference database comprising the *RdRp* sequences of HNU5 and HNU6. Using BLASTN alignment, a total of 3,609 matching sequences were identified. Following quality control measures, which included the removal of sequences shorter than 300 bp, a final dataset of 2,314 *RdRp* sequences with varying lengths was obtained.

Subsequently, γ-CoV sequences (excluding IBV strains) were filtered to retain a single representative sequence per host species with >95% nucleotide identity. This filtering process, combined with the inclusion of 10 geographically diverse IBV sequences, a total of 153 partial γ-CoV *RdRp* sequences were obtained for phylogenetic tree construction. For comparative purposes, we also analyzed an unfiltered dataset containing 611 non-IBV sequences, with the corresponding phylogenetic tree presented in Figure S1.

Phylogenetic trees were constructed using IQ-tree v2.1.3 [29] and Fast Tree [30], and the best-fitting nucleotide substitution model was calculated by the ModelFinder according to the Bayesian information criterion (BIC) method [31].

### 2.5 Recombination analysis

The full-length genomic sequences of γ-CoV were aligned using MAFFT v7.149 [28], and subsequently subjected to systematic recombination analysis through RDP5 software, which implements seven distinct detection methods (RDP, GENECONV, Bootscan, Maximum chi square, Chimera, SISCAN, and 3SEQ) [32]. To ensure the reliability of the results, recombination events were considered statistically significant only when concurrently identified by at least three independent methods. Furthermore, we implemented the Bonferroni correction method with a significance threshold of p≤0.05 to minimize false positive rates. For validation, the identified recombination breakpoints were confirmed through comparative sequence analysis using Simplot v3.5.1, examining the aligned sequences of recombinants, major/minor parental strains, and background sequences.

### 2.6 Coevolutionary analysis

Paired evolutionary trees of γ-CoVs and their corresponding hosts were used to examine the evolutionary relationship. γ-CoVs strains without host information and completely *RdRp* gene were excluded in this analysis. Then, for γ-CoVs from the same host and the identity is greater than 95%, one is randomly reserved as a representative. The host evolutionary tree and topology were obtained from the TimeTree website through entering the Latin text of all species to be analyzed [33]. Phylogenetic tree of virus was constructed by IQ-tree v2.1.3 [29].

Based on the former research of RNA virus evolution [34], we used the eMPRess program [35] to investigate the cospeciations (co-divergence) and cross-host transfer events between γ-CoVs and their hosts. Consistency test of paired host–virus tree in eMPRess was performed on the basis of a duplication-transfer-loss (DTL) model, using a maximum parsimony approach (MPR) to find a “best” mapping of the virus tree onto the host tree which minimized the total “cost.” The null hypothesis was that the similarity of the “true” host-virus tree is not greater than the similarity of host-virus trees generated randomly. The “cost” of every event was set as follows: cospeciations = 0 (fixed), duplication = 1, transfer (also named host switch) = 1, and loss = 1 [36]. The statistical test of co-divergence was carried out by comparing the estimated cost to null distributions calculated from 100 random permutations of the host tip mappings.

### 2.7 Estimation of divergence dates

Following the retrieval of all full-length γ-CoV sequences from GenBank, regression of root-to-tip distances using Treetime software [37] indicated a poor temporal signal among full-length sequences and sampling time (Figure S5). Consequently, node calibration, rather than tip calibration (sampling time), was employed for divergence time estimation in this study.

Given the lack of comprehensive γ-CoV divergence time references, we established calibration points based on well-characterized δ-CoV evolutionary timelines [7]. We implemented a stringent filtering protocol for the γ-CoV *RdRp* dataset (excluding IBV), retaining only one representative sequence per unique combination of collection year, host species, and 100% sequence identity. This dataset and representative IBV sequences yielded a final set of 148 *RdRp* sequences from γ-CoV and δ-CoV for molecular clock analysis. Then, we tested the best combination of molecular clocks and tree priors model using the path sampling, and the uncorrelated relaxed clock with the constant size model as the optimal configuration, based on maximum marginal likelihood value (Table S5).

The *RdRp* sequences were aligned using the MAFFT v7.149 program in BioAider v1.527 [27, 28]. ModelFinder was then utilized to determine the optimal substitution model, which was identified as GTR+F+I+G4 [31]. Next, BEAST v1.10.4 was employed to estimate divergence times, with a Markov chain of 50 million steps and samples taken every 10,000 steps[38]. Divergence time calibrations (ya, years ago, SD, Stdev) were based on previous studies, with normal distribution parameters as follows: HKU11 (mean of 106 ya and SD of 50 ya), HKU17/HKU30 (mean of 319 ya and SD of 50 ya), HKU16-related lineage (mean of 824 ya and SD of 50 ya), and MT138104/ MT138105/ MT138108 (mean of 50 ya and SD of 50 ya) with normal distribution[7]. The prior mean evolution rate was 2.0 × 10^−4^ subs/site/year (1.0 × 10^−4^–1.0 × 10^−3^, 95% highest posterior density or HPD) with uniform distribution [7], and the time to the most recent common ancestor (tMRCA) was calculated using an uncorrelated lognormal relaxed clock. Tracer v1.7 was used to assess the Effective Sample Size (ESS) for all parameters, ensuring convergence by confirming ESS values above 200. The Maximum Clade Credibility (MCC) tree was generated by Tree Annotator v1.10.4 after discarding the initial 10% of states.

## 3 Results

### 3.1 γ-CoVs in wild bird

A total of 252 wild-bird fecal samples were collected in Kunming, and 230 in Dali, covering fourteen bird species in total. After RT-PCR screening with specific primers (Table S1), twelve γ-CoV positive samples were detected, resulting in a prevalence of 2.49%. Specifically, three γ-CoV positive samples were identified in Black-headed gulls (*Chroicocephalus ridibundus*) from Kunming, while the remaining nine samples were collected from Dali: three samples from Common shelducks (*Tadorna tadorna*), two from Gadwalls (*Mareca strepera)*, two from Mallards (*Anas platyrhynchos*), one from a Greylag goose (*Anser anser*), and one from a Ruddy shelduck (*Tadorna ferruginea*) (Table 1). Five out of six host species belong to the order Anseriformes, indicating host specificity.

**TABLE 1.**
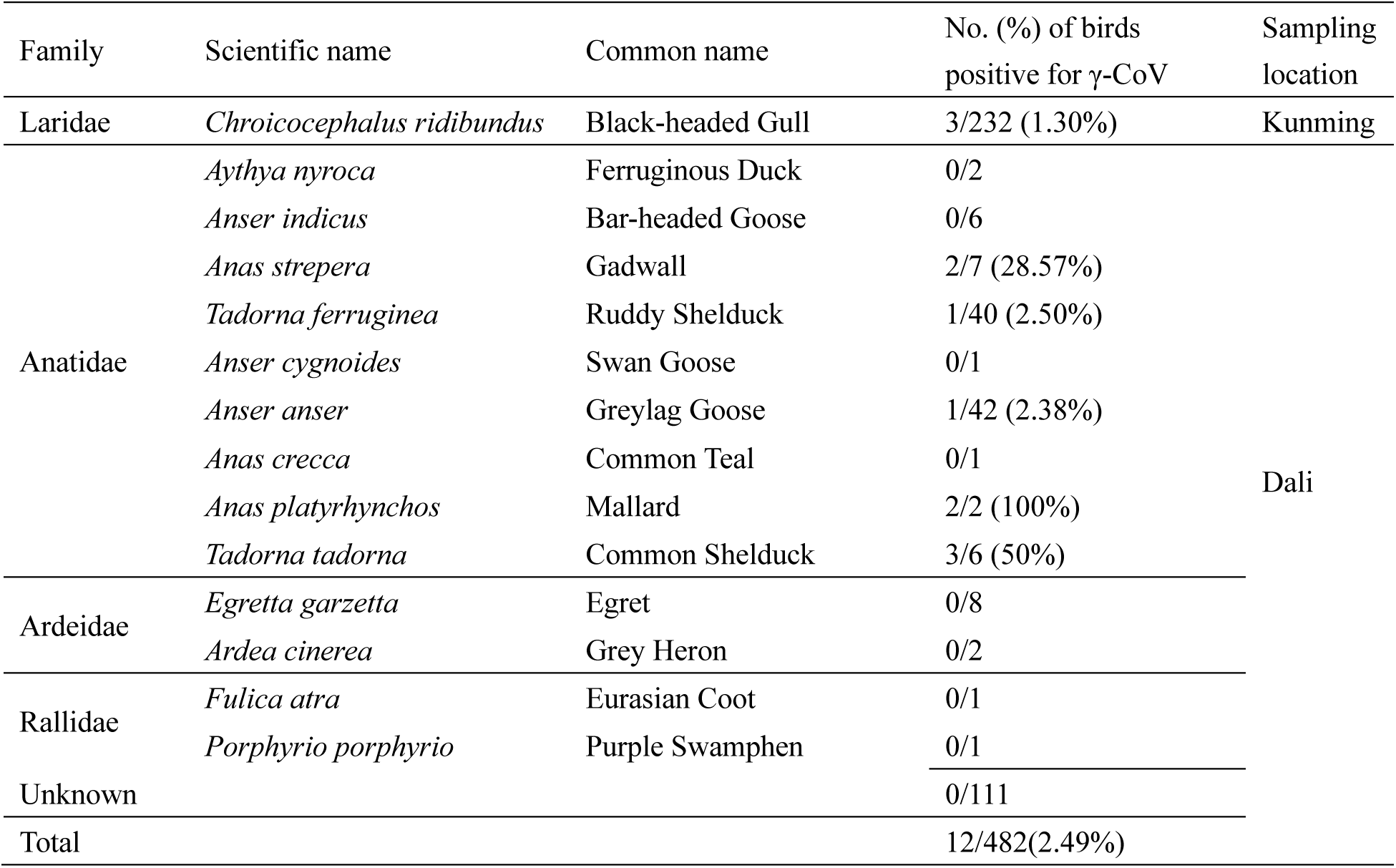
Prevalence of γ-CoV in wild-birds.

### 3.2 Genomic characterization

We successfully obtained complete genome sequences of five γ-CoVs, named HNU5-1, HNU5-2, HNU5-3, HNU6-1, and HNU6-2, using a set of RT-PCR amplifications, Sanger sequencing, and assembly (Table S3). For positive samples with low virus loads, we amplified partial genome sequences (Table1). To preliminarily confirm the taxonomy of these positive samples, we constructed a phylogenetic tree with other genera of CoVs using about 350bp *RdRp* sequences. The result showed that the sequence identified in this study clustered within the subgenus of *Igacorovirus* of γ-CoV (Figure 1, S1).

**Figure 1.**
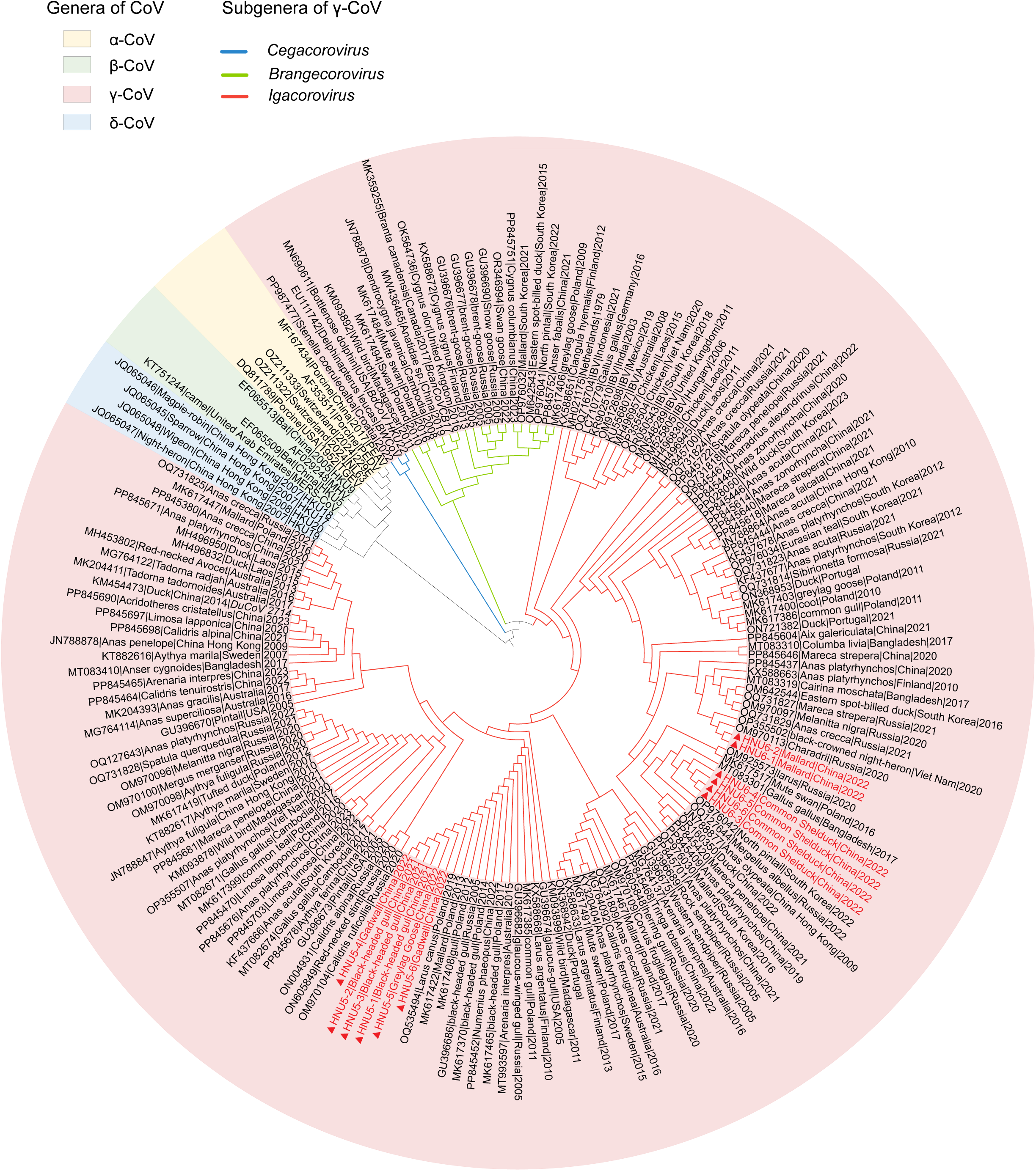
Phylogenetic analyses of partial *RdRp* gene of based on nucleotides. The tree was constructed by using TIM2+F+I+G4 as the substitution model. The background color represented different genera of CoV: α-CoV was indicated by yellow, β-CoV by green, δ-CoV by blue, and γ-CoV by pink. The color of branch in γ-CoV represented different subgenera: *Cegacorovirus* in blue, *Brangecorovirus* in green, and *Igacorovirus* in red. The label marked in red ones were sequences found in this study.

For HNU5-1/2/3 (HNU5s), genomic sequence lengths ranged from 27,305 to 27,874 nt, and the average G + C content varied between 39.54% and 39.68%. The strains of HNU5 possessed a classic genomic structure: 5′ untranslated regions (UTR)-ORF1ab, spike (S), envelope (E), membrane (M), and nucleocapsid (N)-3′UTR. The additional ORFs are diverse, and they were named in the order on the genome. HNU5s were predicted to encode ORF4a, ORF4b, ORF5a, and ORF5b, which are located between M and N. (Figure 2, Table S2). According to the identity analysis, HNU5s share more than 97% sequence identity. The closest relative to HNU5 was a γ-CoV strain (GenBank accession: PP845452) isolated from *Numenius phaeopus* in Shanghai, China, sharing 95.45% whole-genome sequence identity. While the E and M genes showed more than 95% identity, the S gene exhibited 93% identity, and the N gene displayed lower identity at 89%, suggesting potential recombination events in these genomic regions. For HNU5-4/5/6, we amplified partial *RdRp* sequences with fragment lengths of 405 nt, 396 nt, and 342 nt, respectively. The identity between HNU5-4/5/6 and HNU5s was 97.70%, based on the overlapping 340nt fragment.

**Figure 2.**
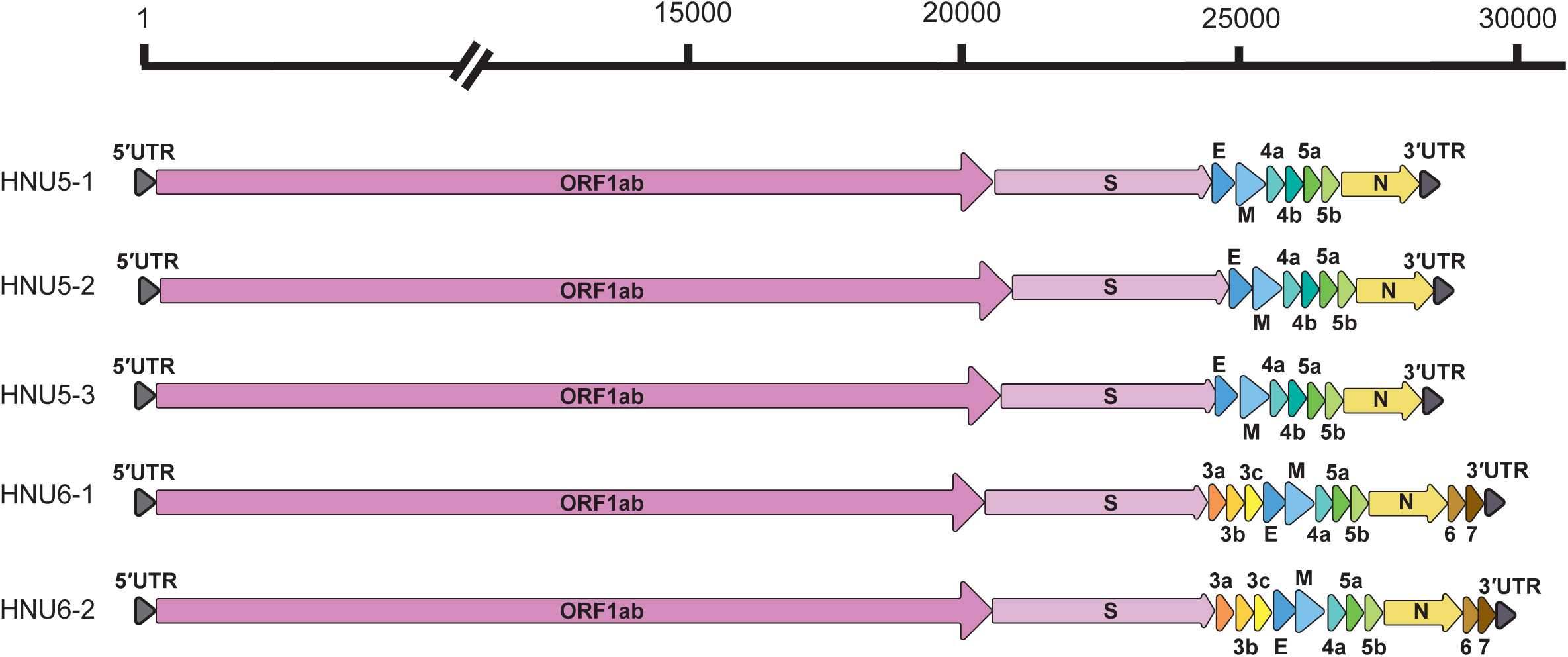
Genomic structure of five γ-CoVs. Each genome included the 5 ′ UTR, ORF1ab, S, E, M, N, and various accessory proteins (e.g., 3a, 3b, 3c, 4a, 4b, 5a, 5b, 6, and 7). The 3′UTR was also shown. Genomic lengths (in base pairs) were indicated, illustrating the variability among the genomes.

For HNU6-1/2, genomic sequence lengths were 29,622 and 29,745 nt, and the average G + C content were 39.8% (Table S2). In addition to the typical ORFs, their additional ORFs, 3a, 3b, and 3c, were located sequentially downstream of the S gene and upstream of the E gene. The ORFs of 4a, 5a, and 5b were located between M and N, while ORF6 and ORF7 were located downstream of N (Figure 2). The HNU6-1/2 strains showed the highest similarity to an γ-CoV strain (GenBank accession: PP845437) isolated from *Anas platyrhynchos* in Shanghai, China. Furthermore, more than 95% amino acid identity across all structural proteins, further supporting their close phylogenetic relationship (Table S3). For HNU6-3/4/5/6, we amplified partial *RdRp* sequences with fragment lengths of 1392 bp, 1671 bp, 1611 bp, and 1698 bp, respectively. After aligning the six HNU6s sequences, the overlapping region of approximately 1400 bp was selected to calculate the identity, which was approximately 97% between HNU6-3/4/5/6 and HNU6-1/2.

The replicase polyprotein ORF1ab occupies more than two-thirds of the entire genome of these five γ-CoVs, and is cleaved into fifteen non-structural proteins (nsps) by nsp3 (putative papain-like protease, PLpro) and nsp5 (putative chymotrypsin-like protease, 3CLpro). Five nsps serve as the remarkable hallmarks for classifying the CoVs, which were 3CLpro, nidovirus RdRp-associated nucleotidyl transferase domain (NiRAN), *RdRp*, zinc-binding domain (ZBD), and superfamily 1 helicase domain (HEL1) [36]. According to species classification in CoVs, a virus can be considered a single species when the sequence divergence across the five domains mentioned above is greater than 7.6%. To confirm the classification of these γ-CoVs, we compared the amino acid sequences of these five domains with those of known γ-CoVs (Table 2). The identity of the five concatenated conserved domains indicates that HNU6-1/2 may belong to the DuCoV_2714 species within *Igacovirus* subgenus. However, HNU5s, with 91.04% identity to DuCoV_2714 in conserved domains, may represent a new species. The results of phylogenetic tree construction based on these five domains and complete genome sequences of γ-CoV also revealed that HNU5s and HNU6-1/2 belong to the *Igacovirus* subgenus (Figure S2, FigureS3).

**Table 2.** Comparison of amino acid identities (%) by combining five conserved replicase domains (3CLpro, NIRAN, *RdRP*, ZBD and HEL) of γ-CoVs.

|  | DuCoV_2714 | BcanCoV_CB17 | BWCoV | ACoV_9203 | IBV | HKU22 |
| --- | --- | --- | --- | --- | --- | --- |
| HNU6-1 | 92.68 | 81.7 | 67.78 | 80.14 | 85.11 | 67.97 |
| HNU6-2 | 92.68 | 81.7 | 67.78 | 80.14 | 85.11 | 67.97 |
| HNU5-1 | 91.04 | 82.02 | 67.72 | 79.57 | 84.67 | 67.91 |
| HNU5-2 | 91.04 | 82.02 | 67.72 | 79.57 | 84.67 | 67.91 |
| HNU5-3 | 91.04 | 82.02 | 67.72 | 79.57 | 84.67 | 67.91 |

Transcription-Regulating Sequences (TRS) are indeed key components of CoV genome, primarily involved in the transcription process. We also utilized Corsid to identify TRS motifs in the CoV genomes found in this study. Two core sequences of TRS were identified as follows: 5′-AAAACGG -3′ and 5′-AACAAA -3′. The latter sequence was found in HNU6-1/2 and aligns with classical TRS motifs previously identified in γ-CoVs (Table 3). The presence of the former sequence in HNU5s suggests a potential diversity in γ-CoV TRS motifs.

**TABLE 3.**
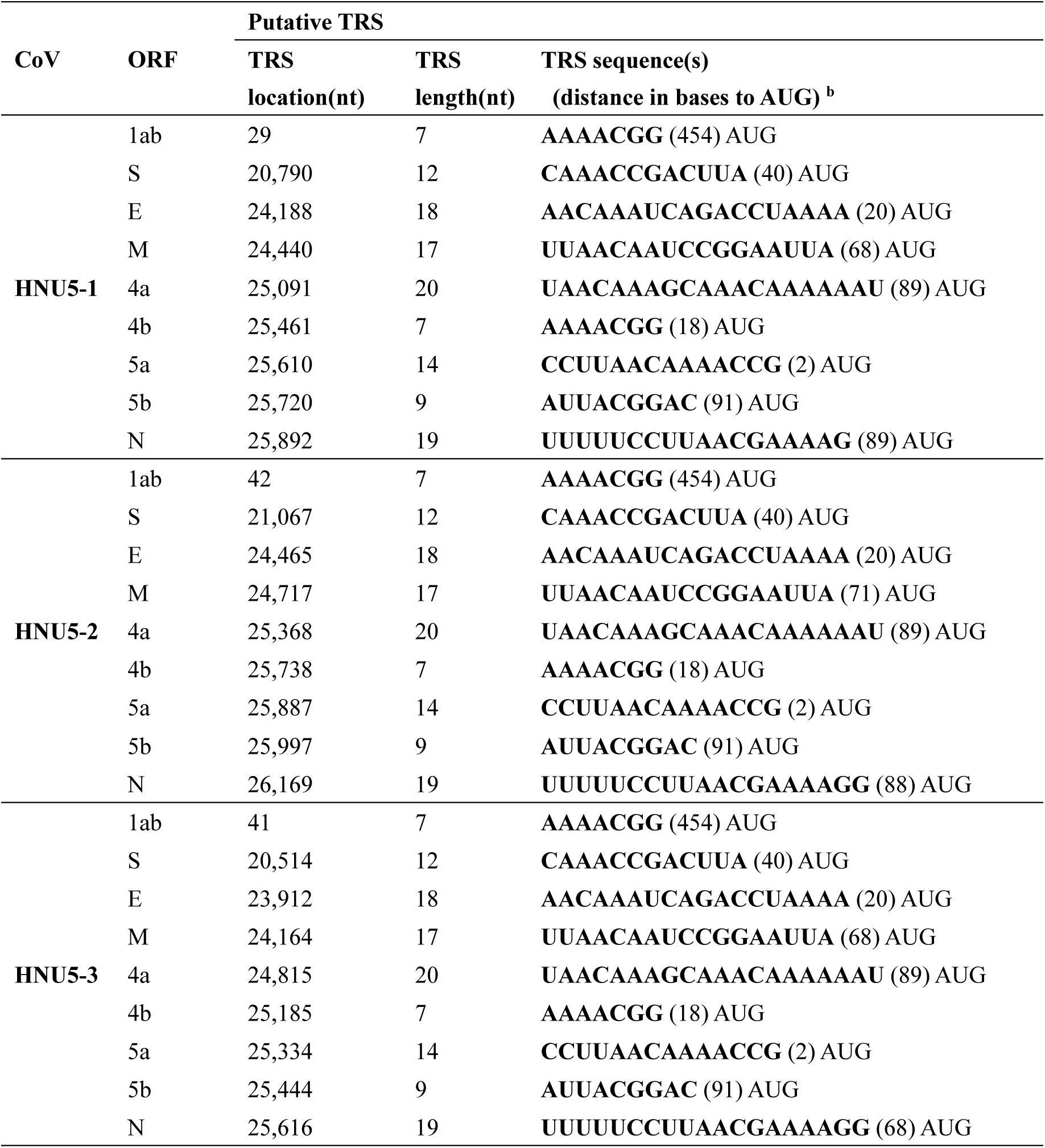
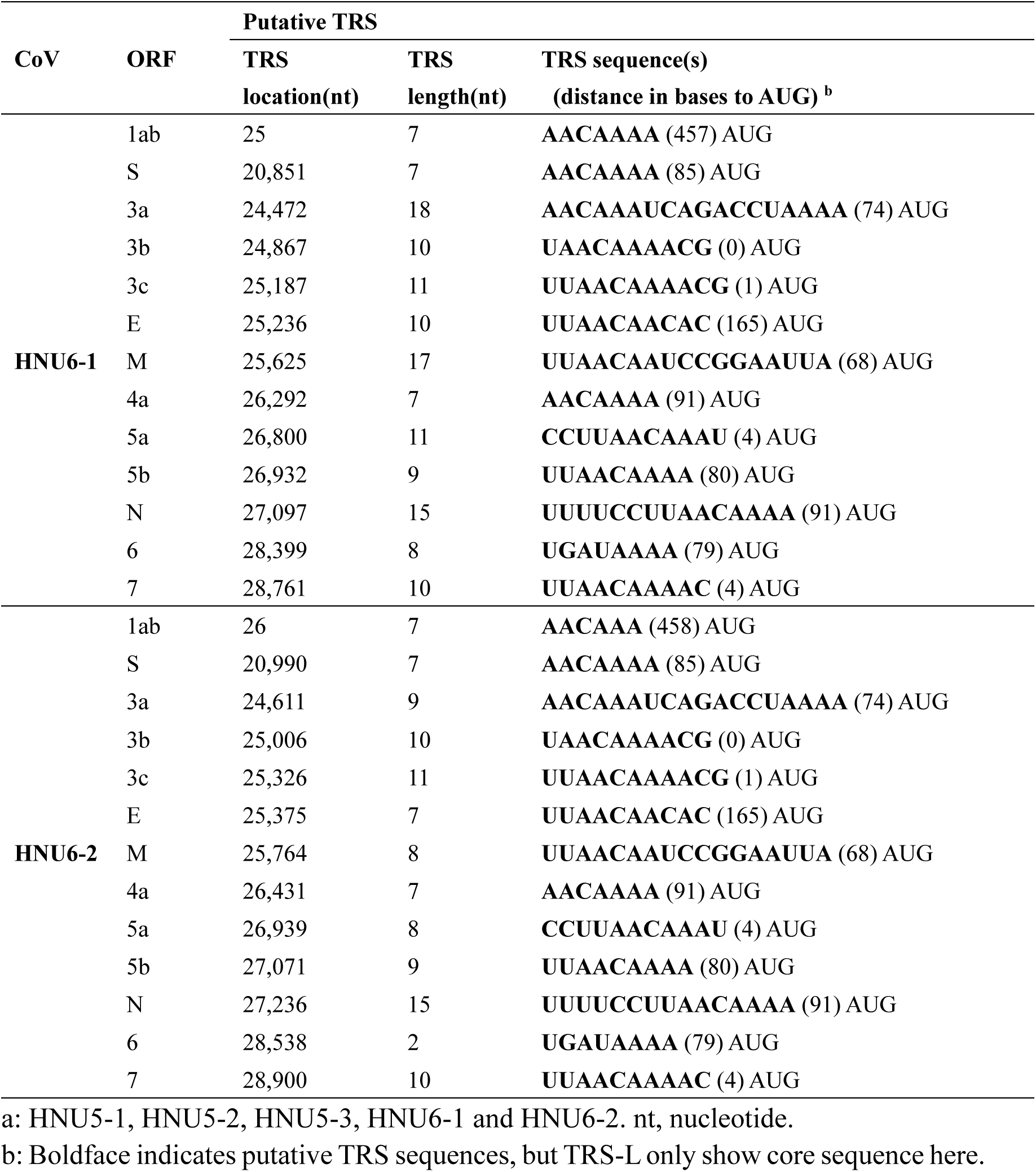
Putative transcription regulatory sequences of CoV genomes ^a^.

### 3.3 Phylogenetics of γ-CoVs

To reveal the evolutionary relationships and genetic conservation of γ-CoVs, we constructed phylogenetic trees based on the amino acid sequences of major genes: ORF1ab, S, E, M, and N.

Phylogenetic analysis revealed that the γ-CoVs identified in this study predominantly clustered within duck-associated lineages, distinct from both IBV-related and marine mammal-associated γ-CoVs. For the S, E, and M genes, it was HNU6-1/2 grouped with MK204411(isolated from *Tadorna tadornoides* in Australia, 2017), and PP845437 (from *Anas platyrhynchos*). In contrast, HNU6-1/2 formed a distinct branch with PP845437 for the N gene, separate from the MK204411. Similarly, HNU5 strains clustered closely with PP845452 (from *Numenius phaeopus*) and OQ535494 (from *Larus canus* in Poland, 2019), consistent with sequence identity analyses. These findings underscore the remarkable genetic diversity within the γ-CoV genus (Figure 3).

**Figure 3.**
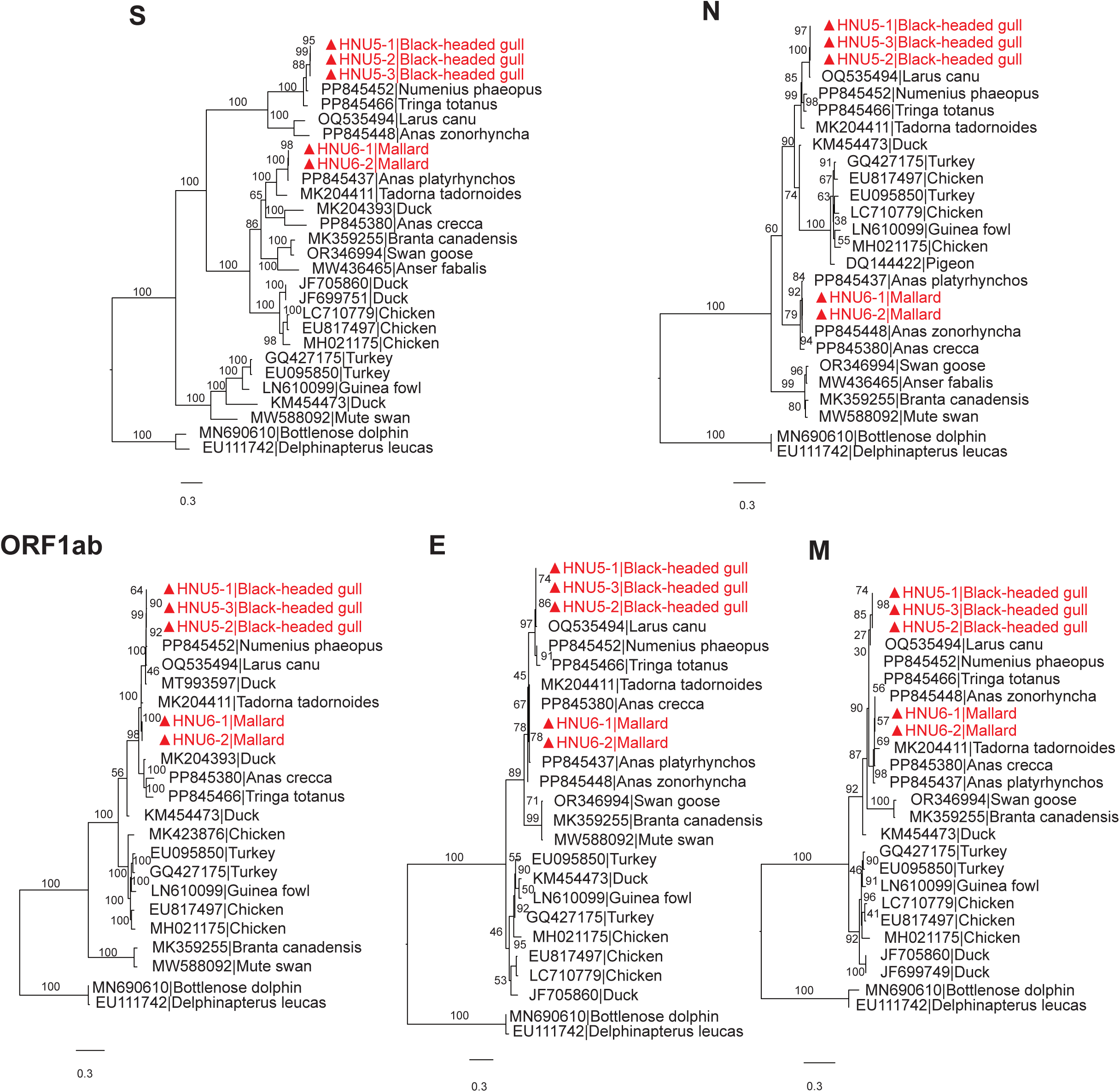
Phylogenetic analyses based on amino acid sequences of ORF1ab, S, E, M and N of γ-CoVs. The substitution model were, in order, JTT+F+G4, WAG+F+G4, Q.plant+G4, LG+G4, and Q.pfam+F+G4. The label on the branch represented the bootstrap value. The marked red ones were γ-CoVs found in this study.

### 3.4 Recombination events of γ-CoV

Putative recombination events among the γ-CoVs were identified using the RDP5 v5.23 software suite [32], incorporating the seven detection methods mentioned above. A highly significant recombination event was detected between PP845448 (major parent, from *Anas zonorhyncha*) and PP845466 (minor parent, from *Tringa totanus*), resulting in the recombinant strain HNU5s. This event was robustly supported by all detection methods, with p-values ≤ 0.05 (Table S4). The breakpoints were precisely identified in RDP5, located at genome positions 20,642–27,351 nt (22,553–35,365 nt in the alignment), corresponding to the spike (S) gene.

Additionally, HNU6 was identified as a recombinant strain originating from PP845380 (major parent, from *Anas crecca*) and HNU5s (minor parent,from *Chroicocephalus ridibundus*). This recombination event was also strongly supported by all methods, with p-values ≤ 0.05. The breakpoints were mapped in RDP5 at genome positions 214–12,911 nt (217–14,686 nt in the alignment), located within the ORF1ab gene (Figure 4). These findings underscore the significant role of recombination in driving the genetic diversity of γ-CoVs.

**Figure 4.**
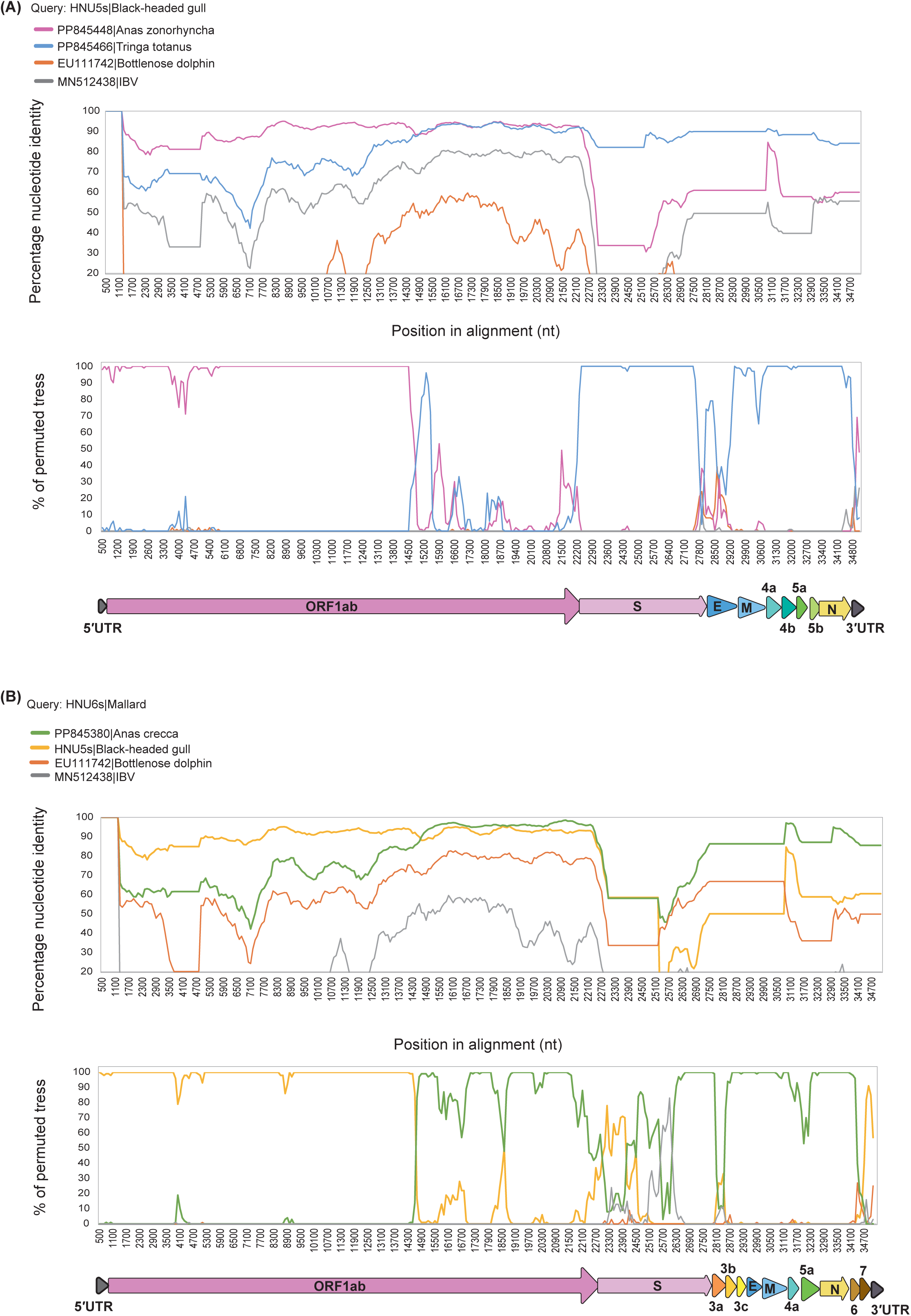
Identification of potential recombination events in γ-CoVs. (A) Potential recombination events detected in the HNU5 strain. (B) Potential recombination events detected in the HNU6 strain. Both the Similarity Plot and Bootscan methods in Simplot v3.5.1 were utilized, with a window size of 1,000 base pairs, and a step size of 100 base pairs. Bootscan analysis was conducted with 1,000 bootstrap replicates for robust validation.

### 3.5 Host-Switching Events of γ-CoV

In the co-divergence and host-switching test, a total of thirty-six events of virus-host evolution were detected: nine co-divergence, six duplications, twenty-one host-switches. The global co-divergence test indicated a significant difference in “cost” between the original and a randomly arranged host-virus tree (p-value < 0.01) (Figure S4), suggesting that γ-CoVs and their hosts generally exhibit a trend of co-evolution. However, some significant cross-species transmission events indicate active host-switching in γ-CoVs (Figure 5).

**Figure 5.**
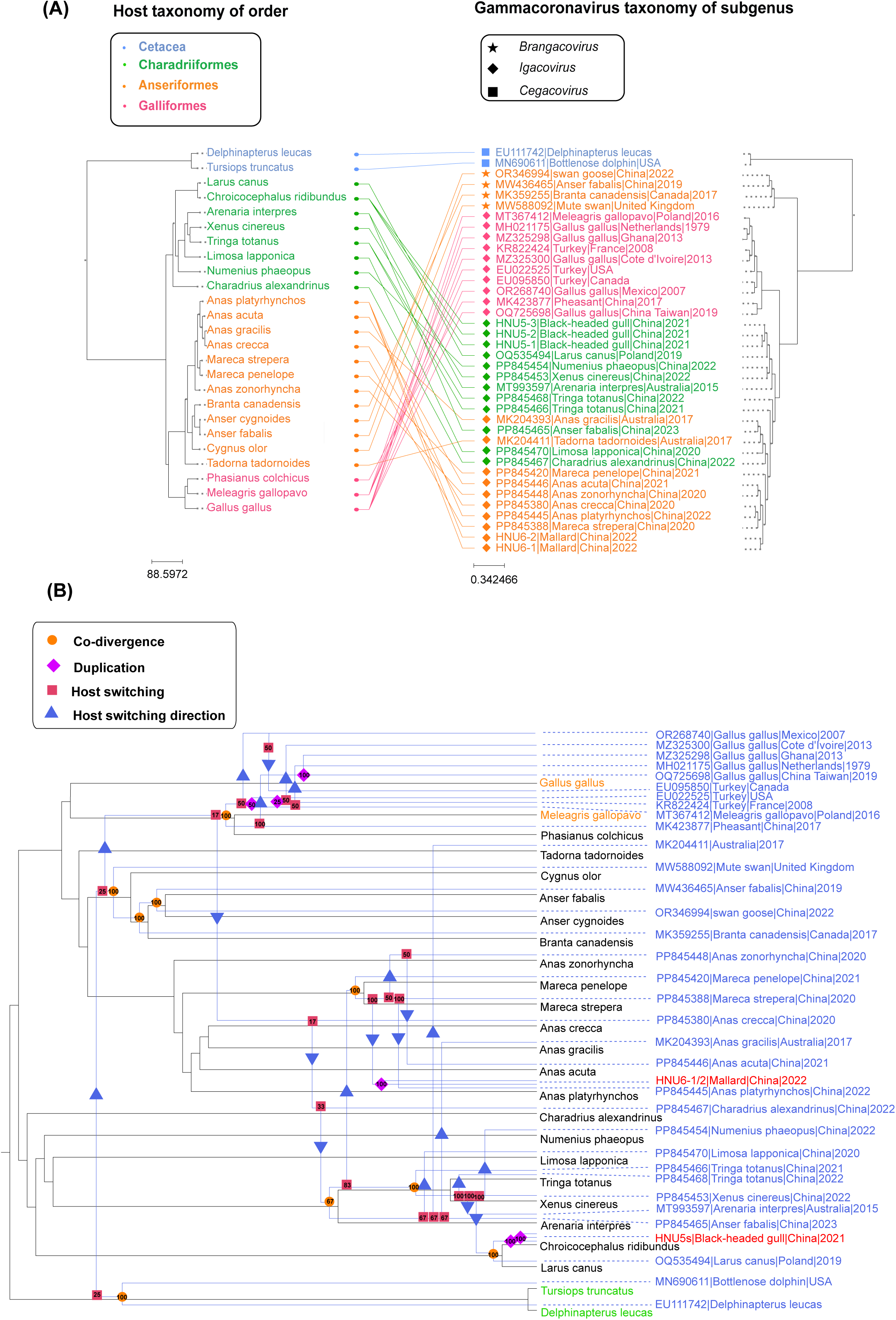
Coevolutionary analysis of virus and host. (A) The tanglegram between host and virus tree. On the left was the host evolutionary tree, and the same color represented the same order. The right was the phylogenetic tree of viral based on the *RdRp* gene, and the same shape represented the same virus subgenus. The tanglegram of host-virus was drawn by PhyloSuite software. (B) Co-divergence and host switch of γ-CoV and their hosts. The host tree was highlighted in black with γ-CoV tree in blue, and each event was labeled with different symbols. Co-divergence and duplication were marked with filled circles and diamonds. Host switching and its direction were marked with squares and arrows, respectively. Losses were not detected in this study. The number above symbols represented the event frequencies. In γ-CoV tree, the marked red ones were γ-CoVs found in this study. In the host tree, the marked orange ones were poultry, the marked black ones were wild-birds, and the marked green ones were marine mammals.

Generally, there were three types of hosts in γ-CoVs based on the host habitat: poultry, wild-birds, and marine mammals. Through evolutionary processes, γ-CoVs appear to have been transmitted among homologous hosts, such as poultry and wild-birds. Notably, γ-CoVs may have also spread from the ancestors of marine mammals to the ancestors of partial Anseriformes in wild-birds, with this transmission event occurring at a frequency of twenty-five. Furthermore, evidence suggests that γ-CoVs could have been transmitted from the ancestors of Galliformes to wild-birds, with an event frequency of seventeen, highlighting the potential for cross-species transmission from poultry to wild-birds. Interestingly, the reverse transmission may also have occurred, as the ancestors of some Anseriformes appear to have transmitted the virus to the ancestors of Galliformes, with an event frequency of twenty-five, indicating that wild-birds could have served as a source of viral transmission to poultry (Figure 5).

In summary, γ-CoVs were found to transmit from marine mammals to wild birds, from wild birds to poultry, from poultry to wild birds, and also involve inter-wild bird, , and inter-poultry transmission. These findings highlighted the complex and directional nature of γ-CoV transmission dynamics among three kinds of hosts.

### 3.6 Divergence Dates of γ-CoV

We estimated the divergence dates of of γ-CoV including HNU5s and HNU6-1/2 by analyzing the *RdRp* nucleotide sequences (Figure 6, S6). The mean evolutionary rate of γ-CoVs was approximately 2.17 × 10^−4^ subs/site/year (95% HPDs, 1.49 × 10^−4^–2.95 × 10^−4^). Using molecular clock dating with the *RdRp* gene, we determined that the tMRCA of δ-CoV and γ-CoV was around 6264 B.C (11946 B.C–2155 B.C, 95% HPD), while the tMRCA of γ-CoV was estimated to be around 1464 B.C (4118 B.C–489 B.C, 95% HPD). The tMRCA of HNU5s was approximately estimated to be 2021 A.D (2016 A.D–2024 A.D, 95% HPD), HNU5 lineage may have diverged from another gull CoV(OQ535494) between 2013 A.D and 2021 A.D. The tMRCA of HNU6-1/2 was approximately 2022 A.D (2017 A.D–2024 A.D, 95% HPD) and HNU6 lineage may have diverged from other duck-related lineages between 1897 A.D and 2022 A.D.

**Figure 6.**
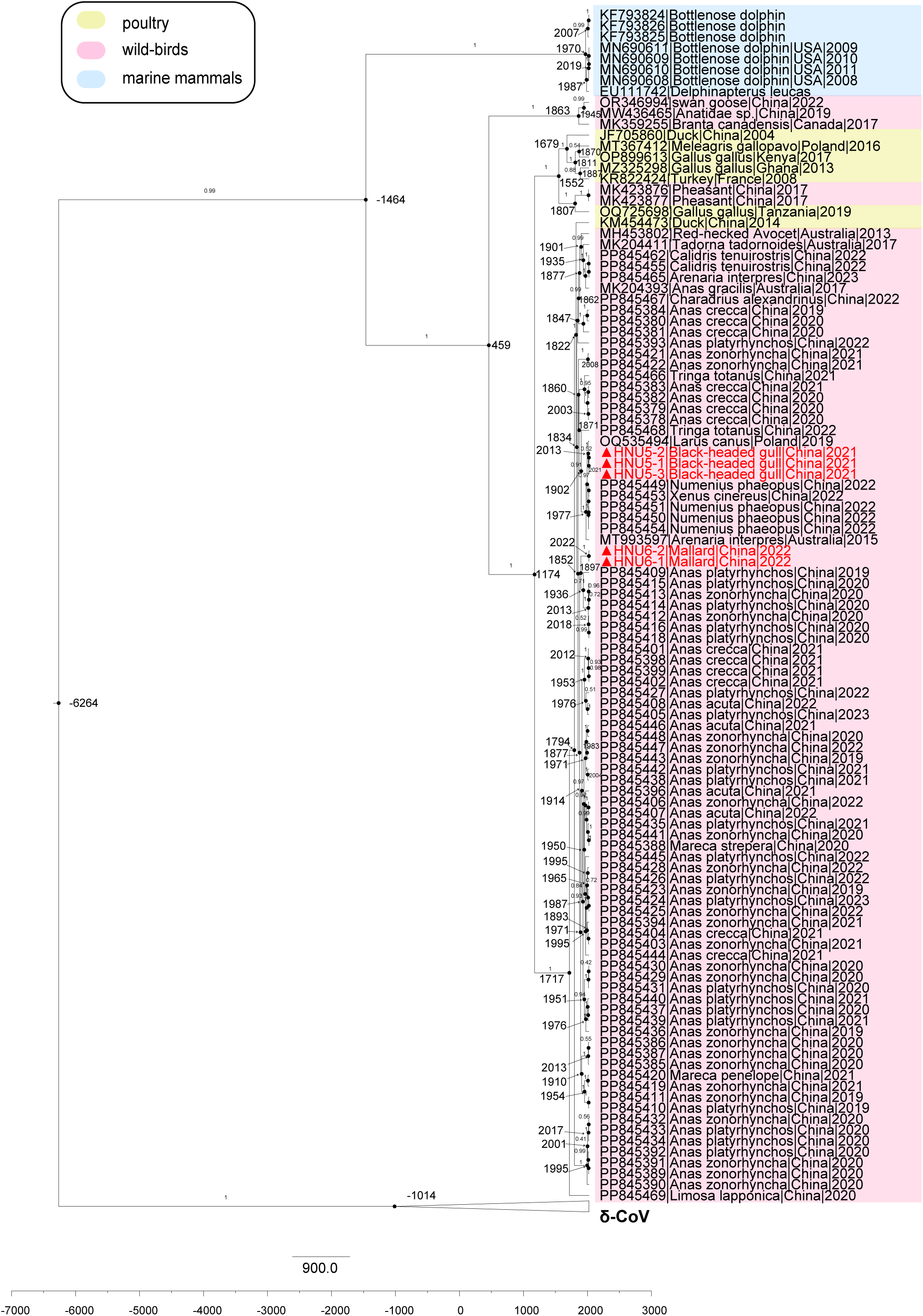
Divergence time based on *RdRp* gene. The value near the node indicated the age of the node, and the label on the branch represented the Bayesian posterior probability. The δ-CoV was collapsed and background colors indicate the host origin: pink represented wild-birds, yellow represented poultry, and blue represented marine mammals. The marked red ones were γ-CoVs found in this study.

## 4 Discussion

Monitoring viruses in wild birds is essential for preventing epidemics and tracing virus transmission. Wild birds, particularly migratory species, can facilitate the spread of viruses and the emergence of cross-regional diseases, such as West Nile Virus and Lyme Disease [18, 39]. In this study, we identified twelve γ-CoVs from 482 wild-bird fecal samples in Yunnan. The positive samples belonged to the class Aves and were categorized into two orders, Charadriiformes and Anseriformes. We amplified and characterized five complete genomes of CoV, designated as HNU5-1, HNU5-2, HNU5-3, HNU6-1, and HNU6-2. Genome comparison revealed that the HNU6-1/2 belong to the DuCoV_2714 species under *Igacovirus* subgenus, while HNU5 genomes represent a new species within the same subgenus. This suggests that γ-CoVs in gulls may differ from other γ-CoVs.

In terms of genomic composition, both strains contained the classic components (5′UTR, ORF1ab, S, E, M, N, and 3′UTR) along with additional ORFs: 4a, 5a, and 5b. However, the amino acid sequence identity within these additional ORFs was relatively low, at approximately 75%, 46%, and 56%, respectively. Additionally, HNU6-1 and HNU6-2 possessed unique ORFs: 3a, 3b, 3c, 6, and 7, while HNU5 genomes possessed ORF4b. Consequently, the designation of these non-classic ORFs was based on their order within the genome rather than their taxonomic classification. Phylogenetic and pairwise identity analyses revealed that HNU5 genomes share close evolutionary relationships with γ-CoV strains from *Larus canus* in Poland (OQ535494) and *Numenius phaeopus* in Shanghai, China (PP845452), despite difference in the S gene sequence. Similarly, HNU6-1/2 strains demonstrated significant S gene variability compared to their close relative, the *Tadorna tadornoides*-associated γ-CoV from Australia (MK204411), highlighting potential recombination events or host-specific adaptation in the spike protein region. These differences in the S gene suggest variations in receptor utilization, potentially enhancing the ability of the virus to transmit across different species and thereby broadening its host range.

Recombination may provide viruses with more evolutionary options than mutations alone [40, 41]. Recombination between CoV occurs frequently and was one of the driving forces behind host-switching. It is associated with CoV spillover, especially in the S gene [42]. The S gene is a key determinant of host species with nearly all major human CoV showing evidence of S gene recombination in their evolutionary history [43]. This recombination can lead to changes in receptor utilization, thereby affecting viral entry. For example, evidence of recombination in the S gene of MERS-CoV suggests that such events are responsible for shifts in receptor binding phenotypes, granting the virus the ability to use human DPP4 as a receptor [44]. Similarly, during recombination events in *Igacoviruses*, the S gene is often fully transferred to another strain, resulting in recombinant viruses [45]. However, data on recombination in γ-CoVs is limited. Avian infectious bronchitis virus (IBV) has been shown to undergo recombination across its entire genome [46], and waterbird γ-CoVs are considered to have originated from recombination with an undescribed γ-CoV strain [47]. In this study, the HNU5 recombinant strain exhibits recombination across the entire S gene region, suggesting a similar mechanism that may influence its host range and receptor binding characteristics. Compared to previous studies, our results provide additional evidence for the high recombination frequency in γ-CoVs, particularly within the S gene. Nevertheless, the extent to which these recombination events influence viral pathogenicity and transmission remains unclear, warranting further investigation.

TRS are responsible for regulating the synthesis of subgenomic mRNAs, which are necessary for the production of viral proteins. TRS is divided into two types: TRS-L (leader), located in the 5’UTR region, and TRS-B (body), located immediately upstream of each viral gene. These elements regulate the transcription of downstream genes [48]. CoV replication is accomplished by the replication and transcription complex, of which *RdRp* is the core component [49, 50]. Sequential synthesis by *RdRp* accomplishes replication and transcription of ORF1a and ORF1b, while discontinuous synthesis enables transcription of other viral genes [48, 51]. This discontinuous transcription is achieved through template switching [52]. During transcription, when *RdRp* hits TRS-B at the viral genome’s 3’ end, it halts and switches to TRS-L, creating negative-strand subgenomic messenger RNAs/ (-) sgmRNAs with a shared leader sequence for each gene. These (-) sgmRNAs serve as templates for (+) sgmRNA synthesis, leading to the production of structural and accessory proteins [52–56]. Both TRS-L and TRS-B contain a conserved core sequence of seven-eight nucleotides (nt), and TRS varies among different genera of CoVs [57, 58]. RNA viruses recombine through template switching, which generates recombinant RNA derived from the genome of two CoVs. CoV transcription involves template switching at TRS-B [59, 60]. Using Corsid, we predicted a distinct TRS sequence in HNU5s: 5’-AAAACGG-3’. Thus, it is possible that these strains may recombine with other CoVs, and this hypothesis has been confirmed in this study.

Due to birds’ flocking behavior and flight capabilities, birds have the potential to transport viruses to other animals and humans, leading to inter-species or cross-species transmission events [5]. Such cross-species transmission may increase the likelihood of emerging zoonotic CoVs [3, 61]. Previous researches have reported avian-to-avian and avian-to-swine transmission caused by δ-CoVs [5]. Within bird populations, frequent host-switching events are occurring due to infections by δ-CoVs [7, 36]. An example of cross-species transmission is HKU15, the only known δ-CoV to infect mammals, which may transmit via sparrows to pigs [36].

However, prior to our study, while frequent transmissions of γ-CoVs within duck species and among various bird species had been documented, there were no reports specifically addressing cross-species transmission between poultry, wild birds, and marine mammals. Based on our coevolutionary analysis, we identified potential host-switching events in γ-CoVs. It appears that γ-CoVs select hosts during their evolution. Our results indicated that γ-CoVs may jump from wild birds to poultry, likely due to the proximity of habitats and similar feeding habits. Habitat loss caused by human activities, leading to winter migratory birds foraging alongside poultry in rice paddies. For example, cross-species transmission of pathogens has been observed between wild red-crowned cranes and domestic geese [62]. It is worth noting that host-switching events have also been observed in the transition from marine mammals to wild birds, possibly due to habitat overlap during migratory periods. However, the event support value was 25%, indicating this clade appeared in 25% of re-samplings, indicating low support so that more evidence is needed to substantiate this cross-species event. Furthermore, transmission events among poultry could lead to significant economic losses. Additionally, γ-CoVs might impact the survival of migratory birds along their migration routes, and they could mutate within wild bird populations, producing new viral strains that pose a greater threat to the farming industry and public health. Although no cases of γ-CoV infection in humans have been reported to date, this study highlights the frequent cross-species transmission risks, particularly the potential routes of transmission from wild-birds to poultry. Furthermore, two newly γ-CoV strains identified in this study are recombinant, underscoring the robust recombination capability of γ-CoV. Therefore, there is an urgent need to enhance surveillance of γ-CoV in animals and to conduct in-depth research on its receptor-binding properties in order to mitigate the potential threat of zoonotic diseases.

In summary, our study suggested the widespread of γ-CoVs among wild-birds and discovered two γ-CoV, HNU5 and HNU6. We have proposed the classification of a new species within the subgenus DuCoV_2714 and new TRS motif in γ-CoVs, which provides new insights to explore γ-CoVs. We have also detected initial evidence of cross-species transmission events among γ-CoVs. This discovery underscores the dynamic nature of coronavirus evolution and the potential for these viruses to jump between different host species. Our findings highlight the importance of continued surveillance and research into the genetic diversity of γ-CoVs to better understand their transmission patterns and inform public health strategies.

## Acknowledgments

This research was jointly funded by the National Natural Science Foundation of China (No. U2002218), the Science and Technology Innovation Program of Hunan Province (2024RC1028), and funds from Hunan University (No. 521119400156).

## Data Availability

The datasets analyzed during the current study are available from the corresponding authors upon reasonable request. All the sequences in this manuscript can be obtained from the NCBI database GenBank with the following accession numbers: from PQ536699 to PQ536710 (https://www.ncbi.nlm.nih.gov, accessed on 31 Oct 2024).

