## Supplementary material for "Sustained cross-species transmission of gammacoronavirus in wild birds reveled by viral characterization in China": Figure S1

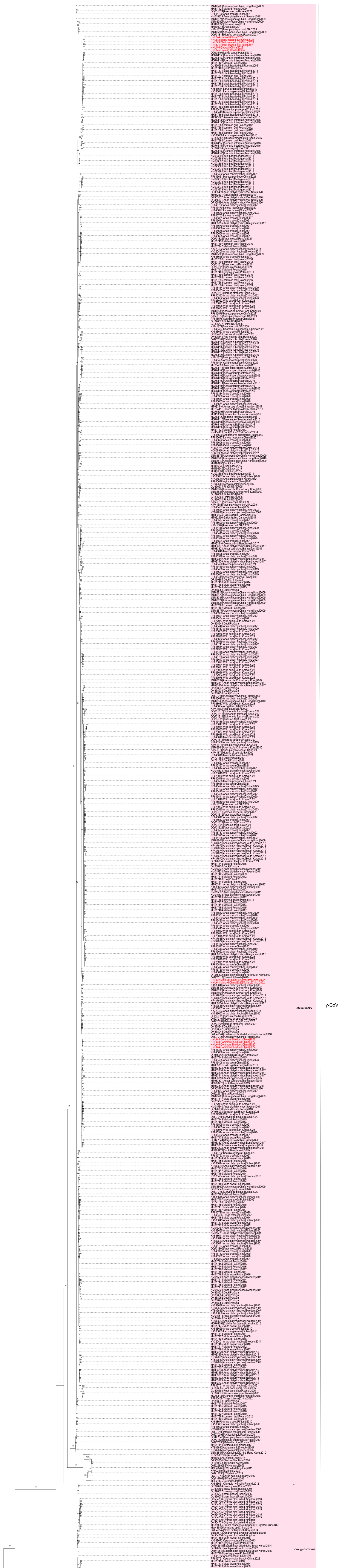


**Figure S1**. Phylogenetic analyses of partial *RdRp* gene of based on nucleotides. The tree was constructed by using TIM2+F+I+G4 as the substitution model. The label on the branch represented the bootstrap value. The color of the branches represented different genera of CoV: α-CoV was indicated by yellow, β-CoV by green, δ-CoV by blue, and γ-CoV by pink.The marked in red ones were sequences found in this study
