## Supplementary material for "Sustained cross-species transmission of gammacoronavirus in wild birds reveled by viral characterization in China": Figure S2

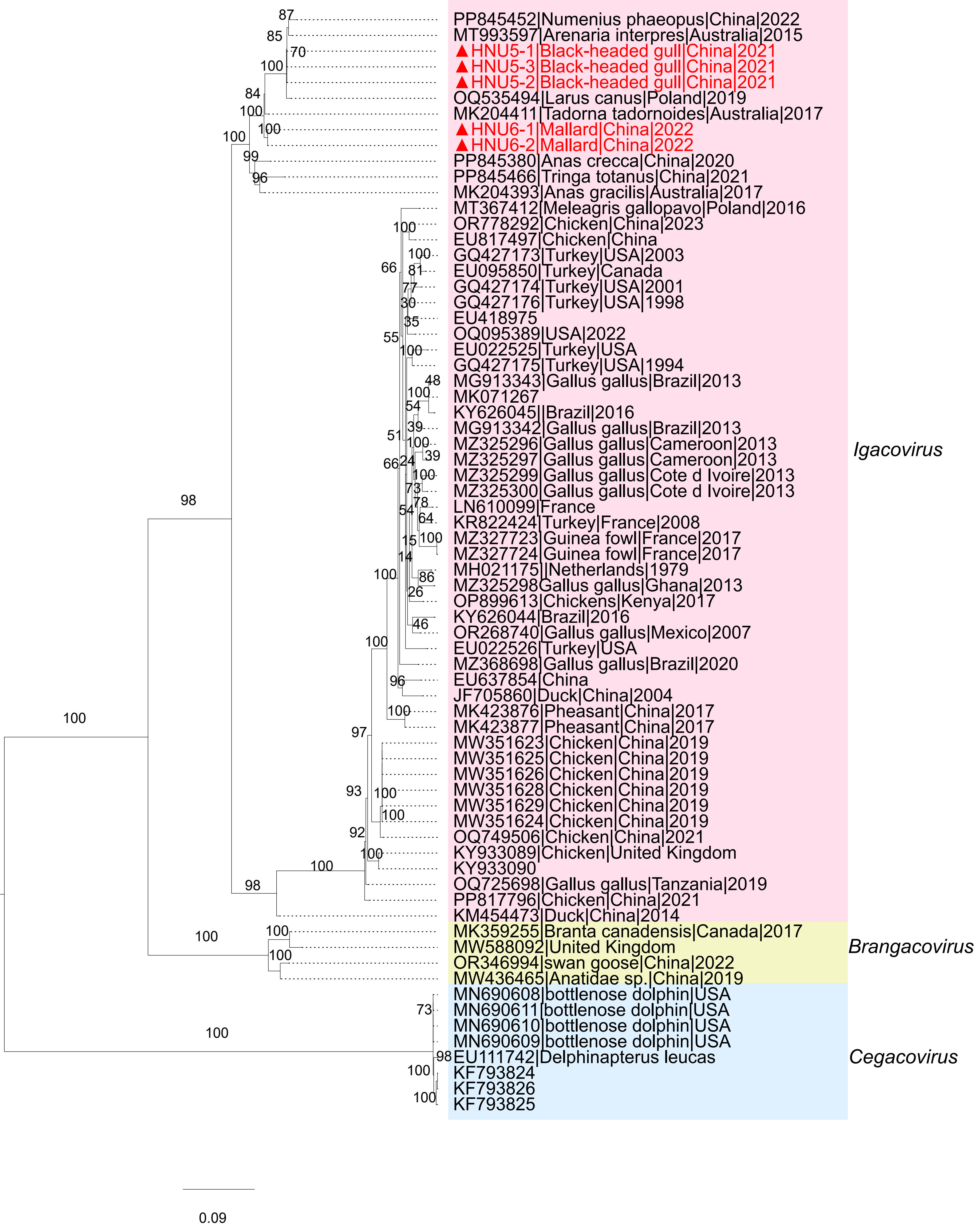


**Figure S2**. Maximum likelihood phylogenetic tree of based on combined amino acids sequences of conserved domains: 3CL-pro, NIRAN, RdRp, ZBD and HEL1. The substitution model was JTT+F+I+G4. The label on the branch represented the bootstrap value. Background colors represented subgenera of CoV: pink represented *Igacovirus*, yellow represented *Brangacovirus*, and blue represented *Cegacovirus*. Red markers indicated the γ-CoVs identified in this study.
