## Supplementary material for "Sustained cross-species transmission of gammacoronavirus in wild birds reveled by viral characterization in China": Figure S3

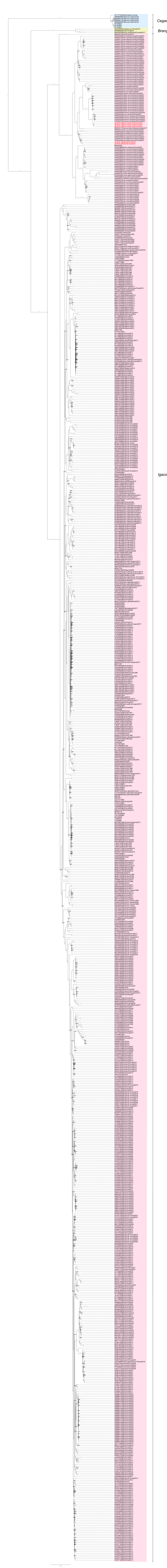


**Figure S3.** Maximum likelihood phylogenetic tree based on complete genome of γ-CoVs. Background colors represented subgenera of CoV: pink represented *Igacovirus*, yellow represented *Brangacovirus*, and blue represented *Cegacovirus*. Red markers indicated the γ-CoVs identified in this study.
