## Supplementary material for "Sustained cross-species transmission of gammacoronavirus in wild birds reveled by viral characterization in China": Figure S4

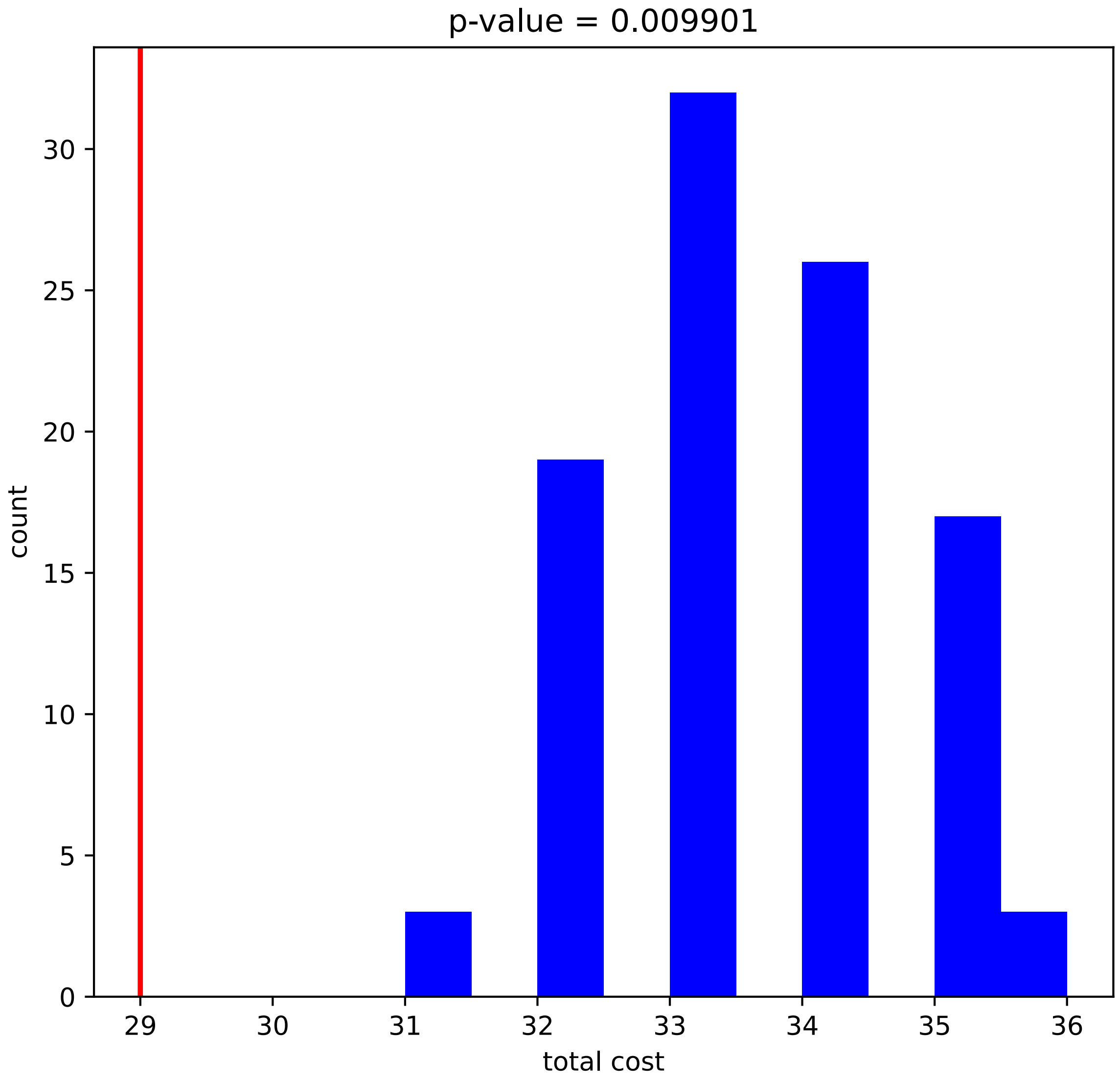


**Figure S4.** The distribution of costs of random sample. The histogram represented the costs random sample, and the cost of original data was marked with a red line.
