## Supplementary material for "Sustained cross-species transmission of gammacoronavirus in wild birds reveled by viral characterization in China": Figure S5

(A)

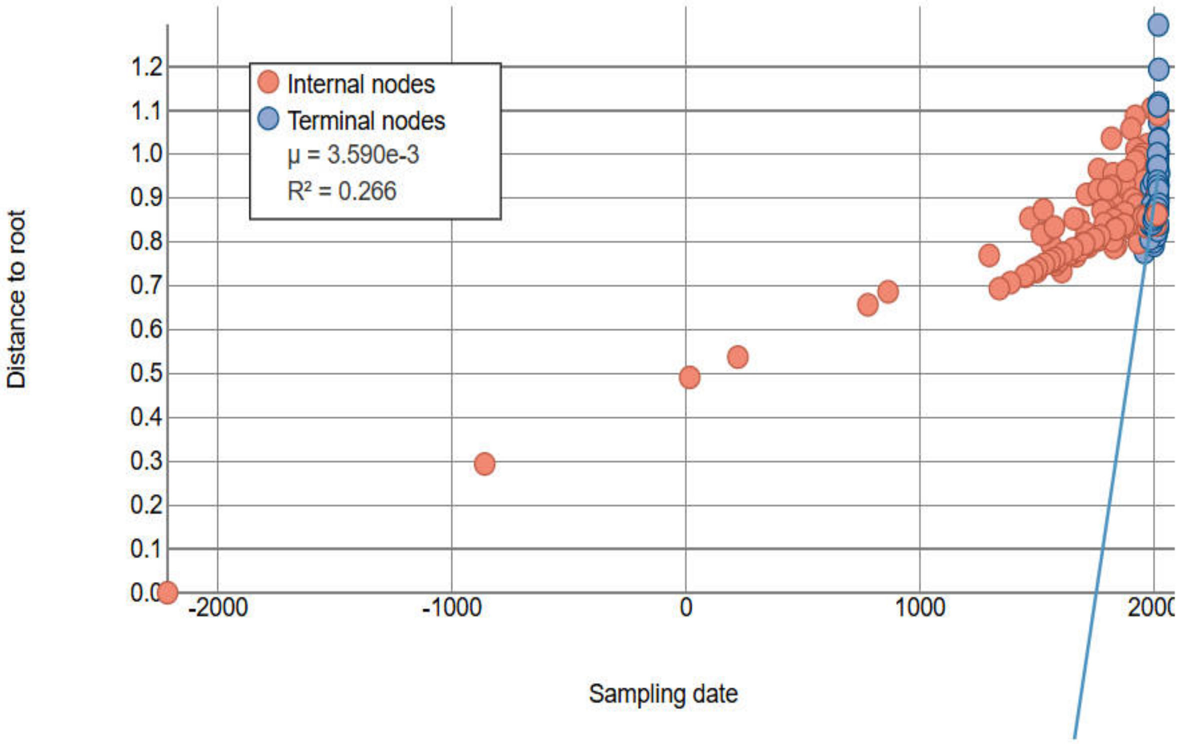


(B)

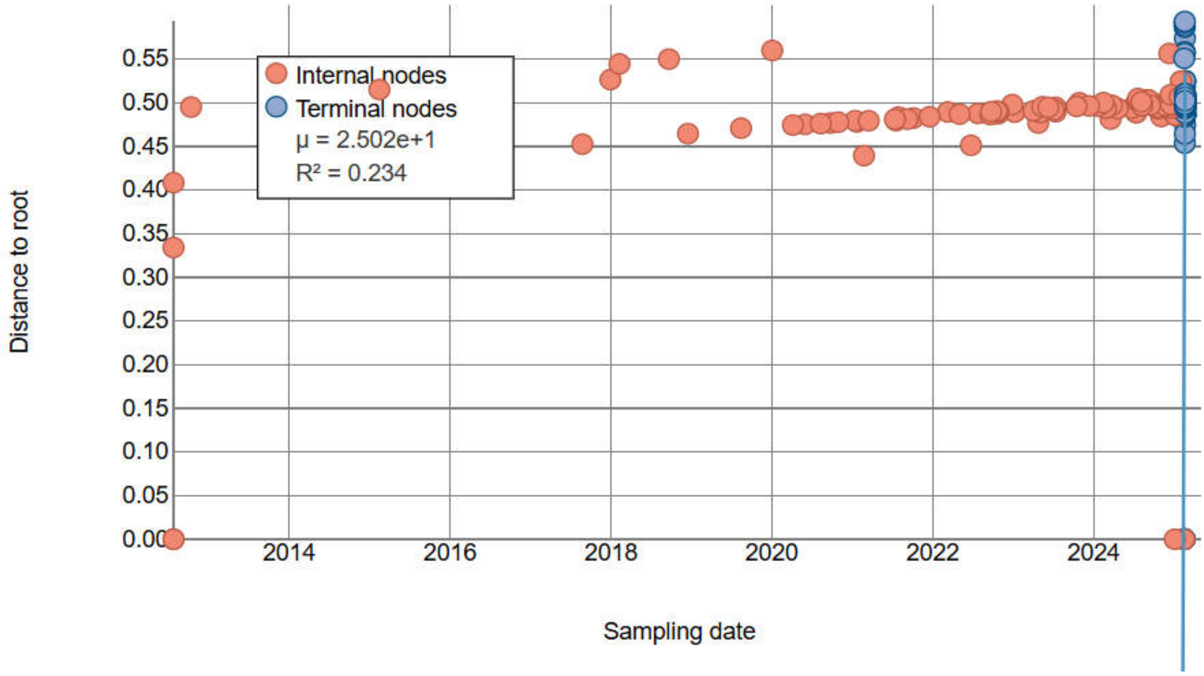


**Figure S5**. Regression analysis of root-to-tip genetic distances. The maximum likelihood (ML) phylogenetic tree was constructed based on the complete genome sequences (A) and RdRp gene (B) of γ-CoV. The *μ* value represented the evolution-ary rate, and the $R^2$ value indicated the correlation coefficient.
