## Supplementary material for "Sustained cross-species transmission of gammacoronavirus in wild birds reveled by viral characterization in China": Figure S6

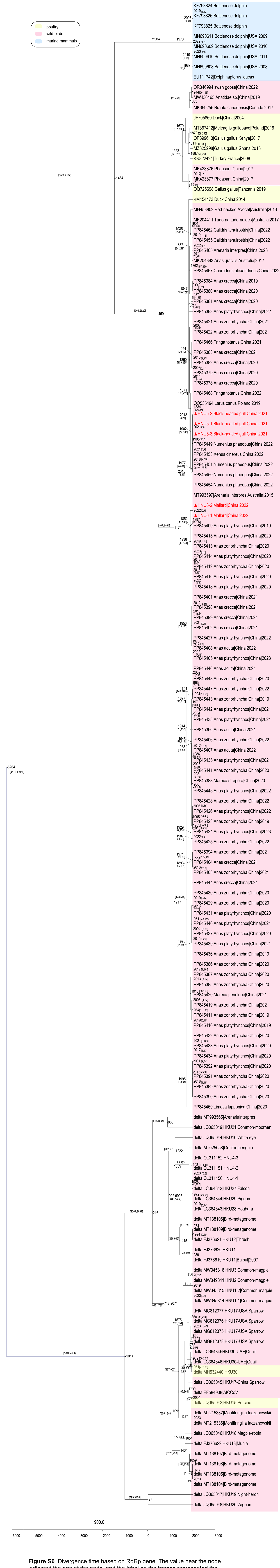

**Figure S6.** Divergence time based on RdRp gene. The value near the node indicated the age of the node, and the label on the branch represented the 95% HPD. Background colors indicated the host origin: pink represented wild -birds, yellow represented poultry and blue represented marine mammals. The marked red ones were γ-CoVs found in this study.
