## Supplementary material for "Sustained cross-species transmission of gammacoronavirus in wild birds reveled by viral characterization in China": Table S1

| **CoV** | **Genome length** | **G + C content (%)** | **ORF** | **Location (nt)** | **Length (nt)** | **Length (aa)** | **Frame** |
| --- | --- | --- | --- | --- | --- | --- | --- |
| **HNU5-1** | 27,608 | 39.59 | 1ab | 640-20,891  Shifting at 2,816 | 20,252 | 6,750 | +1 |
|  |  |  | S | 20,842-24,216 | 3,375 | 1,124 | +1 |
|  |  |  | E | 24,226-24,528 | 303 | 100 | +1 |
|  |  |  | M | 24,525-25,199 | 675 | 224 | +3 |
|  |  |  | 4a | 25,200-25,484 | 285 | 94 | +3 |
|  |  |  | 4b | 25,486-25,629 | 144 | 47 | +1 |
|  |  |  | 5a | 25,626-25,823 | 198 | 65 | +3 |
|  |  |  | 5b | 25,820-26,065 | 246 | 81 | +2 |
|  |  |  | N | 25,999-27,255 | 1,257 | 418 | +1 |
| **HNU5-2** | 27,874 | 39.68 | 1ab | 503-21,168  Shifting at 12,942 | 20,666 | 6,888 | +1 |
|  |  |  | S | 21,119-24,493 | 3,375 | 1,124 | +2 |
|  |  |  | E | 24,503-24,805 | 303 | 100 | +2 |
|  |  |  | M | 24,802-25,476 | 675 | 224 | +1 |
|  |  |  | 4a | 25,477-25,761 | 285 | 94 | +1 |
|  |  |  | 4b | 25,763-25,906 | 144 | 47 | +1 |
|  |  |  | 5a | 25,903-26,100 | 198 | 65 | +1 |
|  |  |  | 5b | 26,097-26,342 | 246 | 81 | +3 |
|  |  |  | N | 26,276-27,532 | 1,257 | 418 | +2 |
| **HNU5-3** | 27,305 | 39.54 | 1ab | 502-20,615  Shifting at 12,531 | 20,114 | 6,704 | +1 |
|  |  |  | S | 20,566-23,940 | 3,375 | 1,124 | +1 |
|  |  |  | E | 23,950-24,252 | 303 | 100 | +1 |
|  |  |  | M | 24,249-24,923 | 675 | 224 | +3 |
|  |  |  | 4a | 24,924-25,208 | 285 | 94 | +3 |
|  |  |  | 4b | 25,210-25,353 | 144 | 47 | +1 |
|  |  |  | 5a | 25,350-25,547 | 198 | 65 | +3 |
|  |  |  | 5b | 25,544-25,789 | 246 | 81 | +1 |
|  |  |  | N | 25,723-26,979 | 1,257 | 418 | +1 |

**TABLE S2**. Coding potential of five γ-CoV genomes.

| **CoV** | **Genome length** | **G + C content (%)** | **ORF** | **Location (nt)** | **Length (nt)** | **Length (aa)** | **Frame** |
| --- | --- | --- | --- | --- | --- | --- | --- |
| **HNU6-1** | 29,622 | 39.80 | 1ab | 489-20,992  Shifting at 12,918 | 20,504 | 6,835 | +3 |
|  |  |  | S | 20,943-24,500 | 3,558 | 1,185 | +3 |
|  |  |  | 3a | 24,564-24,869 | 306 | 101 | +3 |
|  |  |  | 3b | 24,877-25,182 | 306 | 101 | +1 |
|  |  |  | 3c | 25,199-25,411 | 213 | 70 | +2 |
|  |  |  | E | 25,411-25,713 | 303 | 100 | +1 |
|  |  |  | M | 25,710-26,390 | 681 | 226 | +3 |
|  |  |  | 4a | 26,390-26,674 | 285 | 94 | +2 |
|  |  |  | 5a | 26,815-27,015 | 201 | 66 | +1 |
|  |  |  | 5b | 27,021-27,272 | 252 | 83 | +3 |
|  |  |  | N | 27,203-28,447 | 1,245 | 414 | +2 |
|  |  |  | 6 | 28,486-28,764 | 279 | 92 | +1 |
|  |  |  | 7 | 28,775-29,299 | 525 | 174 | +2 |
| **HNU6-2** | 29,745 | 39.85 | 1ab | 490-21,131  Shifting at 13,056 | 20,642 | 6,881 | +3 |
|  |  |  | S | 21,082-24,639 | 3,558 | 1,185 | +3 |
|  |  |  | 3a | 24,703-25,008 | 306 | 101 | +3 |
|  |  |  | 3b | 25,016-25,321 | 306 | 101 | +1 |
|  |  |  | 3c | 25,338-25,550 | 213 | 70 | +2 |
|  |  |  | E | 25,550-25,852 | 303 | 100 | +1 |
|  |  |  | M | 25,849-26,529 | 681 | 226 | +3 |
|  |  |  | 4a | 26,529-26,813 | 285 | 94 | +2 |
|  |  |  | 5a | 26,954-27,154 | 201 | 66 | +1 |
|  |  |  | 5b | 27,160-27,411 | 252 | 83 | +3 |
|  |  |  | N | 27,342-28,586 | 1,245 | 414 | +2 |
|  |  |  | 6 | 28,625-28,903 | 279 | 92 | +1 |
|  |  |  | 7 | 28,914-29,438 | 525 | 174 | +2 |

* HNU5-1, HNU5-2, HNU5-3, HNU6-1 and HNU6-2. nt, nucleotide; aa. Amino acid.
